# Repetitive magnetic stimuli over the motor cortex impair consolidation of a balance task by suppressing up-regulation of intracortical inhibition

**DOI:** 10.1101/2024.09.12.612606

**Authors:** Sven Egger, Michael Wälchli, Samuel Meyer, Wolfgang Taube

**Author notes:** Corresponding Author: Sven Egger, University of Fribourg, Bd de Pérolles 90, Office F440, 1700 Fribourg/Freiburg, Switzerland.

## Abstract

Low-frequency repetitive transcranial magnetic stimulation (rTMS) over the primary motor cortex (M1) was shown to impair short-term consolidation of a balance task, emphasizing the importance of M1 in balance skill consolidation. However, the disruptive mechanisms of rTMS on neural consolidation processes and their persistence across multiple balance acquisition sessions remain unclear. GABAergic processes are crucial for motor consolidation and, at the same time, are up-regulated when learning balance skills. Therefore, this study investigated the impact of rTMS on GABA-mediated short-interval intracortical inhibition (SICI) and consolidation of balance performance. Participants (n=31) underwent six balance acquisition sessions on a rocker board, each followed by rTMS (n=15) or sham-rTMS (n=16). In the PRE-measurement, SICI was assessed at baseline and after balance acquisition with subsequent rTMS/sham-rTMS. In the POST-measurement, this procedure was repeated to assess the influence of motor memory reactivation on SICI. In addition, SICI-PRE and SICI-POST were compared to assess long-term processes. Both groups achieved similar improvements within the balance acquisition sessions. However, they did not consolidate equally well indicated by significant declines in performance for the rTMS group (*p* = 0.006) in the subsequent sessions. Both short-(*p* = 0.014) and long-term (*p* = 0.038) adaptations in SICI were affected by rTMS: while the sham-rTMS group up-regulated SICI, rTMS led to reductions in inhibition. The interfering effect of rTMS on both balance consolidation and up-regulation of SICI suggests that increased intracortical inhibition is an important factor to protect and consolidate the newly acquired motor memory.

## Introduction

The primary motor cortex (M1) is thought to play an important role for the acquisition and the consolidation of motor tasks due to its considerable plasticity (Bütefisch et al., 2000; Classen et al., 1998; Della-Maggiore et al., 2015; Karni et al., 1995; Karni et al., 1998; Sanes & Donoghue, 2000). Acquisition of a motor task leads to rapid online performance improvements, which form an unstable motor memory trace (Dayan & Cohen, 2011; Kantak & Winstein, 2012). During consolidation, this newly created motor memory trace undergoes offline (i.e., without further practice) stabilization, enabling long-term storage of the motor memory (McGaugh, 2000; Robertson et al., 2004). However, it is important to note that the motor memory trace is not permanently stable after consolidation. Upon reactivation (i.e., when the motor memory is retrieved), the memory trace may temporarily destabilize, necessitating reconsolidation (Dudai et al., 2015; Walker et al., 2003). A widely applied method to test consolidation (and reactivation) processes and thus plasticity in M1 is transcranial magnetic stimulation (TMS). With this method, it was shown that excitability in M1 is decreased after several weeks of balance training (Beck et al., 2007; Schubert et al., 2008; Taube et al., 2007). At the same time, the level of short-interval intracortical inhibition (SICI) was demonstrated to increase after learning balance tasks in young (Mouthon & Taube, 2019; Taube et al., 2020) and elderly people (Kuhn et al., 2024). This increase in inhibition was assumed to be important for the long-term consolidation process (Bachtiar & Stagg, 2014; Kida & Mitsushima, 2018) and was speculated to contribute to shifting movement control from initially more cortical to more subcortical levels after movement automatization (Mouthon & Taube, 2019).

So far, it is well established that sufficient time is needed to consolidate the acquired motor memory and to protect it from interference (Brashers-Krug et al., 1996). One possibility to induce interference and thus, disrupt motor memory consolidation is to learn a second task shortly after learning a first task. In this way, interference effects have been shown for simple motor tasks, such as force-field adaptations (Brashers-Krug et al., 1996; Shadmehr & Brashers-Krug, 1997), visuomotor rotations (Krakauer et al., 1999; Tong et al., 2002; Wigmore et al., 2002), ballistic force tasks (Lundbye-Jensen et al., 2011; Roig et al., 2014), and sequence learning (Neville & Trempe, 2017). Recently, we have demonstrated such interference effects also for a complex, whole-body task, involving balancing on a rocker board (Egger et al., 2021). Thus, learning a balance task B involving the same muscles as a previously learned balance task A did interfere with the consolidation of the first balance task (A), indicating that the two motor tasks were competing for similar neural resources in the brain. However, with this experimental design, it is impossible to designate specific brain structures that are involved in the consolidation of the task(s). Therefore, induction of more focal interference effects is warranted that allow the identification of the involved brain structures. One way to test more directly the involvement of M1 for learning and consolidation is application of repetitive TMS (rTMS). Repetitive TMS can transiently disrupt the function of M1 and consequently may interfere with M1-dependent motor behavior/learning by creating temporary “virtual lesions” (Siebner et al., 2009). With this non-invasive method, interference was reported after learning tasks that are thought to strongly rely on M1 such as ballistic finger (Baraduc et al., 2004; Muellbacher et al., 2002), ankle (Lundbye-Jensen et al., 2011), and wrist motor tasks (Cothros et al., 2006; Roig et al., 2014). Most importantly with respect to the present study, rTMS was shown to interfere with short-term motor memory consolidation in the balancing task on the rocker board (Egger et al., 2023). In this previous study, participants underwent one single acquisition phase of 48 trials of balancing on the rocker board, followed by either 15 min of low-frequency rTMS or sham-rTMS, and finally a retention test 24 h later. In this retention test, the rTMS group revealed a performance loss while the sham-rTMS group displayed off-line gains. This is strong evidence that the primary motor cortex plays a crucial role in consolidating balance skills.

To date, the exact mechanisms how low-frequency suprathreshold rTMS disrupts the consolidation of motor tasks is poorly understood. It is assumed that rTMS decreases corticospinal excitability (CSE) for certain tasks and thus, may interfere with cortical processes (Censor & Cohen, 2011). It has further been suggested that rTMS is affecting primarily synaptic plasticity (Hoogendam et al., 2010) but inconsistent findings were found when applying paired-pulse paradigms to test for changes in SICI (Chen et al., 2018; Daskalakis et al., 2006) so that review articles concluded that rTMS has no or only minor effects on SICI (Fitzgerald et al., 2006). However, the great majority of these mechanistic studies were done while subjects were at rest although it is well known that neural circuits are activated in a strongly task-specific manner (Opie & Semmler, 2016; Soto et al., 2006). Therefore, the present study tested neural markers of plasticity not only at rest (i.e., upright stable stance) but also while participants were actually executing the previously learned balance task. We hypothesised that rTMS would interfere not only with the consolidation process of the balance skill but also with adaptations in SICI. To test this, two balance learning groups were formed, one receiving rTMS and the other sham-rTMS immediately after each of the 6 balance acquisition sessions. We assumed that with increased automatization of the balance task, the role of M1 would decrease and thus, rTMS-induced interference effects would be less pronounced. Furthermore, rTMS was expected to interfere with the commonly known up-regulation of SICI after balance training (Mouthon & Taube, 2019; Taube et al., 2020).

## Materials & methods

### Participants

In total, 31 healthy young volunteers participated in the study. Participants signed an informed consent form. The study procedures were reviewed by the local ethics committee (Comission cantonale d’éthique de la recherche sur l’être humain (CER-VD); ID: 2022-01526), were in accordance with the declaration of Helsinki, and were declared risk-free considering the relevant TMS exclusion criteria (Rossi, 2009). Participants were randomly allocated into an intervention group (rTMS: *n* = 15, 8 women, 23 ± 2 years, 65 ± 10 kg, 1.75 ± 0.1 m) or a control group (sham-rTMS: *n* = 16, 8 women, 26 ± 6 years, 68 ± 13 kg, 1.74 ± 0.07 m). To reduce variability in TMS measurements as much as possible, PRE- and POST-measurements were done around the same time of the day (Matamala et al., 2018; Sale et al., 2007), and subjects were asked to abstain from confounding substances, such as caffeine or medication (Pellegrini et al., 2020).

### Experimental design

The experimental design (Fig. 1) included the acquisition and consolidation of a balance task for two different intervention groups (rTMS group, sham-rTMS group). Behavioral and neural adaptations in response to balance training were assessed in the short-term (‘Short-term adaptation’ and ‘Reactivation’; Fig. 1, red) and the long-term (Fig. 1, blue). All participants completed the same balance learning task. However, in order to disturb consolidation, one group received rTMS after each balance session (rTMS group) whereas the participants of the sham-rTMS (=control) group received no repetitive magnetic pulses. During the PRE and POST sessions, neurophysiological measurements were conducted before the balance training (PRE_1, POST_1) as well as after rTMS or sham-rTMS (PRE_2, POST_2). Balance acquisition sessions were guided by qualified study personnel. Each session started with a balance familiarization (6 trials) on the rocker board, followed by behavioral (i.e., balance performance) and neurophysiological measurements in the PRE- and POST-sessions. This included the determination of a) the resting motor threshold (rMT) during sitting, b) the active motor threshold (aMT) during slight muscular activity as well as SICI measurements during quiet upright stance (control condition) and during balancing on the rocker board (balance condition). After that, peripheral nerve stimulation (PNS) was applied to assess the maximal M-wave. Subsequently, the learning of the balance task took place, which included 24 trials divided into four series (Series 1(_S1_) to Series 4(_S4_)). Participants rested for 30 s between trials and for 1 min between series. Immediately after the balance acquisition, rTMS or sham-rTMS stimulation was applied. Within both the PRE- and the POST-session, SICI- and PNS-measurements were repeated to assess ‘short-term adaptations’ (PRE_2) as well as the effect of reactivating the previously learned motor task (‘reactivation’; POST_2). In each of the four acquisition sessions, rMT was determined in order to apply rTMS with the correct stimulation intensity (i.e. 115% of rMT). All measurements and acquisition trials were completed either in socks or barefoot.

**Figure 1:**
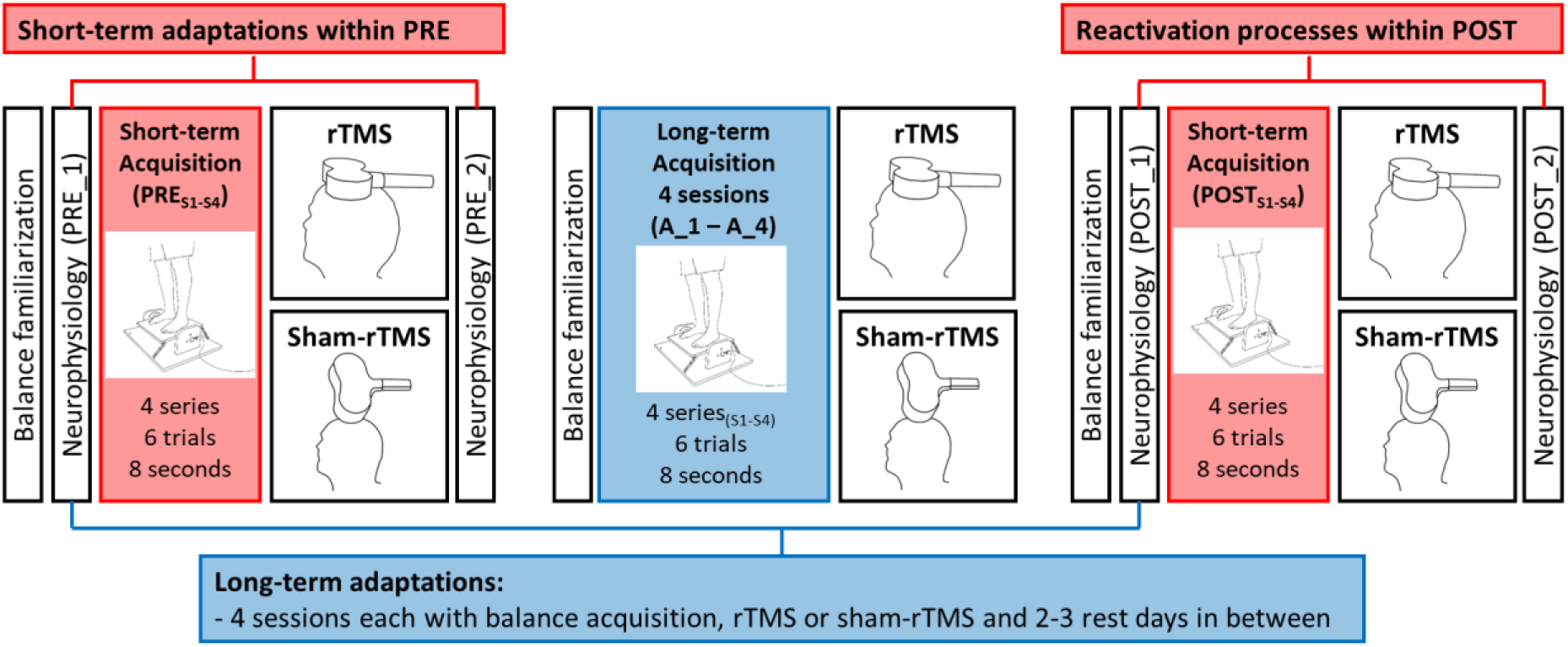
Summary of the study design for the two intervention groups (rTMS: *n* = 15; sham-rTMS: *n* = 16). The colours distinguish between short (red), and long-term (blue) effects. Behavioral short-term adaptations were measured within PRE and POST and behavioral long-term adaptations over 4 balance acquisition sessions. Neurophysiological short-term effects were evaluated twice; in the PRE-measurement, short-term adaptations were assessed PRE_1 to PRE_2. In the POST-measurement, the effects of reactivating the previously learned balance task were investigated from POST_1 to POST_2; ‘reactivation’. Long-term adaptations (i.e. 5 acquisition sessions separated by 2 to 3 days) were assessed from the initial PRE-measurement (PRE_1) to the POST-measurement (POST_1). In the figure, “Neurophysiology” stands for the assessment of corticospinal excitability (single pulses with TMS), SICI (short-interval intracortical inhibition measured by a paired-pulse TMS-paradigm), determination of the resting motor threshold (in order to adjust the relative intensity of the rTMS measurements) and the active motor threshold (in order to adjust the relative intensity of the SICI and corticospinal excitability measurements) and M_max_ (with peripheral nerve stimulation). S1-S4 = series one to series four.

### Balance task and balance learning

A custom-made rocker board (see Fig. 2) was used as a balance device, which allowed forward and backward rotations around a fixed axis (Egger et al., 2021, 2023). Rotations were measured as the angular position of the mediolateral axis by a goniometer (MP20, Megatron Elektronik, Putzbrunn, Germany) with a sampling frequency of 2 kHz. The task difficulty was individualized in the PRE-session during the balance familiarization with different supporting springs (spring constants from 0.124 to 1.119 N/mm) and then maintained for each participant for the entire study. The difficulty was selected so that the subjects scored an initial mean deviation from the rocker board horizontal of around 8° within the 8 s trials. This relatively high challenge was chosen to avoid ceiling effects by guaranteeing that each participant had enough potential to improve their performance throughout the study. Participants were asked to maintain bipedal balance with as little deviation from the horizontal rocker board position as possible. Thus, the aim was to reduce mean deviation to the lowest possible value (i.e., zero). In order to facilitate learning, participants received augmented feedback about the mean deviation in degrees after each trial (Egger et al., 2021, 2023). At the beginning of every trial, participants started with a stable horizontal position on the rocker board by holding on to a laterally positioned handrail. As soon as the hand was detached from the handrail, the trial started. A trial was valid as long as the participant neither left the rocker board nor used the handrail. Otherwise, the trial was repeated.

**Figure 2:**
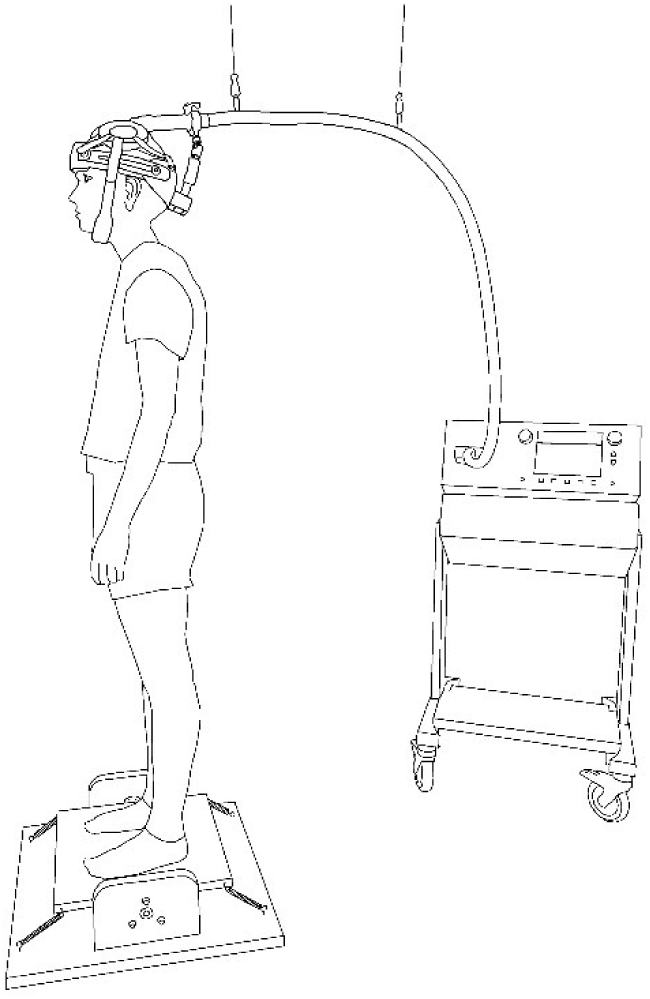
Experimental setup for the transcranial magnetic stimulation with the coil fixture and the custom-made rocker board.

### Transcranial magnetic stimulation (TMS)

Two different figure of eight coils were used for the TMS measurements. Repetitive TMS was performed with a water-cooled double coil (MCF-B70, Tonica Elektronik A/S, Farum, Denmark). Single- and paired-pulse stimulations were applied with an air-cooled double coil (D-B80, Magventure A/S, Farum, Denmark). Both coils were connected to a MagVenture stimulator (MagPro X100 with MagOption MagVenture A/S, Farum, Denmark). The coil was oriented with the handle to the rear and the current passed in an anterior-posterior direction. For each session, the hotspot for the soleus muscle (SOL) was located in a seated position and directly marked with color on the scalp. Subjects were asked not to remove the marker for the duration of the study. Subsequently, rMT was determined in a sitting position with relaxed leg muscles using the MCF-B70 coil. The determination of the resting threshold was necessary to adjust the intensity of rTMS correctly. Afterwards, the aMT was determined during light muscular activity in plantar flexion (monitored by the EMG signal, which should be approximately 0.05 mV during the determination). The aMT was determined with the D-B80 coil for the PRE and POST session. Both motor thresholds were searched first in 5% and then in 1% steps of the stimulator output (Rossini et al., 2015). Thresholds were set where three out of five motor evoked potentials (MEPs) were ≥50 µV for rMT or ≥100 µV for aMT (Lundbye-Jensen et al., 2011; Rossini et al., 2015). Lastly but most importantly, the SICI measurements were performed during two conditions: a) quiet upright stance (control condition) and b) balancing on the rocker board (balancing condition). For these measurements, a helmet fixture to hold the D-B80 coil in place was used (see Fig. 2). During SICI measurements, single-pulse MEPs and paired-pulse MEPs were recorded to assess both CSE and intracortical inhibition. The interstimulus interval for the paired pulse MEPs was 2 ms (Roshan et al., 2003; Vucic et al., 2009). The stimulation intensity of the single- (120%-160% of the aMT) and paired-pulses (65%-85% of the aMT) was adjusted individually for each subject in the PRE_1 measurement and fixed for the subsequent session (Garry & Thomson, 2009). For the measurements in the stable standing position, the rocker board was fixed in horizontal position with wooden blocks so that no movement of the rocker board was possible. Measurements during the balancing task were performed with identical and relatively rigid springs (3.537 N/mm) for all participants in order to ensure sufficient postural stability during the continuous TMS applications and to prevent fatigue. A trigger for the stimulations on the rocker board was released by a custom-made, python-based software program if the rocker board was moved from back to front at a velocity less than 40°/s and the rocker board position was close to the horizontal alignment (−1°). TMS measurements were performed in two series, each containing 20 stimuli, in which single- and paired-pulse stimulations were alternately applied with an interstimulus pause of 4 s. This paired-pulse paradigm was applied during stable standing and during the balancing task on the rocker board.

### Peripheral nerve stimulation (PNS)

The tibial nerve was stimulated using an electrical stimulator (AS 100, ALEA Solutions GmbH, Zurich, Switzerland). The anode (5 x 5 cm) was placed below the patella and the cathode (2 cm diameter) was fixed in the popliteal fossa where the optimal stimulation point was determined. The stimulation duration was 1 ms and the interstimulus interval was 4 s. The stimulation intensity was continuously increased until the M-wave reached a plateau and did not increase further. The maximum value was then defined as M_max_. For PNS application and data collection, a custom-made, python-based software was used, with which stimulation control was manually regulated.

### Repetitive transcranial magnetic stimulation (rTMS)

For the rTMS group, low-frequency magnetic stimulations were applied for 15 min at a frequency of 1 Hz and an intensity of 115% of rMT (Egger et al., 2023; Muellbacher et al., 2002) while subjects lay quietly on a couch. For the sham group, the coil was rotated 90° degrees so that the coil was placed orthogonal to the scalp and no magnetic current reached the brain (Baraduc et al., 2004; Egger et al., 2023). The coil position was fixed in each group by a mounting arm (Magic Arm and Super Clamps, Manfrotto, Cassola, Italy). In both groups, MEPs were visually checked and ensured that rTMS produced MEPs of approximately 0.3 mV in the soleus muscle and sham-rTMS did not produce any muscular responses.

### Electromyography

Two surface electrodes (Ag/AgCI, Ambu Blue Sensor P, Ballerup, Denmark) were attached to the muscle belly of the right SOL to record bipolar EMG. Before attaching the electrodes, the muscle area was shaved, abraded, and disinfected. The EMG electrodes were connected to a transmitter (Myon aktos, Myon AG, Schwarzenberg, Switzerland). EMG signals were amplified analogously (x 200), transmitted wirelessly to the receiver station, and then recorded with a sampling frequency of 2 kHz. The EMG data from the rocker board acquisition sessions and the TMS measurements were recorded using a customized software program (IMAGO Record, Pfitec Biomedical Systems, Endingen, Germany). With the help of this software, the muscle signals (M-waves, MEPs, etc.) could be checked online at any time.

### Data processing

Recorded data were analyzed offline using Matlab (R2021b, The MathWorks, Inc., Natick, MA, USA) and Microsoft Excel (Microsoft 365 MSO, Microsoft Corporation, Redmond, WA, USA). The angular position of the rocker board was low-pass filtered (10 Hz) and the mean deviation from the horizontal alignment was calculated over the 8 s trials. To evaluate balance performance, the average performance per series was calculated, with the best and worst trials in each series deleted to minimize outliers (Egger et al., 2021).

MEPs were filtered and analyzed in the time window between 30 and 80 ms after the stimulation. To normalize single-pulse MEPs, M_max_ (mV) was determined using the highest value from the standing PNS measurements to consider peripheral influences and differences between sessions (Groppa et al., 2012; Rossini et al., 2015). To obtain the single-pulse and paired-pulse MEPs, peak-to-peak amplitudes (mV) were calculated, averaged per condition (standing; balancing) and for each measurement (PRE_1, PRE_2, POST_1, POST_2). Mean normalized single-pulse MEPs were used to determine CSE. SICI average values were calculated as the percentage difference between the conditioned paired-pulse and the unconditioned single-pulse using the formula 100-(DP⁄SP*100), in line with previous studies (Kuhn et al., 2017; Mouthon & Taube, 2019). MEPs were removed from the data analysis (2.6% of all stimulations) if the corresponding background EMG activity before the single- or paired-pulse stimulations was higher than two times the standard deviation of the individual session time point (PRE_1, PRE_2, POST_1, POST_2). For this, background EMG activity for each MEP was assessed using the averaged root mean square value evaluated over a time interval of 100 ms before the stimulation (Mouthon & Taube, 2019).

### Statistical analysis

Jamovi (The jamovi project 2022; jamovi version 2.3, computer software, retrieved from https://www.jamovi.org) was used to conduct all statistical tests. The level of significance was set at *p* ≤ 0.05. All behavioral data were normalized to the respective baseline values (PRE to PRE_S1_, POST to POST_S1_ and balance acquisition to Acquisition_1_S1_). The effect sizes of the significant variance analyses were expressed with the partial eta square (*η^2^p*; small: 0.02; medium: 0.13; large: 0.26) and for the significant *t*-tests with the Cohens-*d* (*d*; small: 0.2; medium: 0.5; large: 0.8). If the sphericity assumptions of the variance analyses were violated, Greenhouse-Geisser adjustments were applied.

#### Balance acquisition

Short-term balance acquisition in PRE and POST measurements was tested with a 2 by 2 mixed-design analysis of variance (ANOVA) with TIME (PRE_S1_; PRE_S4_ or POST_S1_; POST_S4_) as within and GROUP (rTMS; sham-rTMS) as between factor. Long-term balance acquisition was tested with an 8 by 2 ANOVA with TIME (A_1_S1_; A_1_S4_; A_2_S1_; A_2_S4_; A_3_S1_; A_3_S4_; A_4_S1_; A_4_S4_) as within and GROUP (rTMS; sham-rTMS) as between factors. To examine the individual training sessions in more detail, balance performance for each acquisition-session (A_1, A_2, A_3, A_4) was tested with 4 independent 2 by 2 ANOVA’s with TIME (A_1_S1_ to A_1_S4_, A_2_S1_ to A_2_S4_, A_3_S1_ to A_3_S4_ or A_4_S1_ to A_4_S4_) as within and GROUP (rTMS; sham-rTMS) as between factors.

#### Balance consolidation

Consolidation of balance skills was measured as the balance performance at the end of one acquisition session compared to balance performance at the beginning of the subsequent acquisition session and was analyzed with three independent 2 by 2 ANOVA’s with TIME (A_1_S4_ to A_2_S1_, A_2_S4_ to A_3_S1_ or A_3_S4_ to A_4_S1_) as within and GROUP (rTMS; sham-rTMS) as between factors. In case of a significant TIME x GROUP interaction, post-hoc tests with Tukey correction were conducted. Finally, to have an overview about all three consolidation phases together, an unpaired *t*-test was calculated with the mean value for each participant across all consolidation phases indicating individual losses or offline gains between the rTMS and sham-rTMS group. Consolidation was not tested in the PRE- and POST-measurements due to several reasons: neurophysiological assessments directly after rTMS/sham-rTMS while executing the same balance task could have biased the results; either due to application of many magnetic pulses and/or due to the additional exposure to the balance exercises and potential fatigue. In addition, such extensive measurements would have not been tolerated by all participants.

#### Short- and long-term neurophysiological effects of rTMS

Adaptations in SICI and CSE during execution of both conditions (standing and balancing) were compared as delta values in comparison to PRE_1 (Δ). Thus, for SICI and CSE, a 3 by 2 ANOVA was performed for both conditions, with TIME (PRE_2; POST_1 and POST_2) as within and GROUP (rTMS; sham-rTMS) as between factors. The significant main effect of GROUP in SICI during execution of the balance task was followed-up by independent samples Student’s *t*-tests and in case of violation of the normal distribution by Mann-Whitney-*U t*-tests.

## Results

### Balance acquisition

Short-term acquisition: Balance performance improved for both short-term balance acquisition phases before the training (PRE: *F*_1,29_ = 44.81; *p* < 0.001; *η^2^p* = 0.61; see Fig. 3A) and after the training (POST: *F*_1,29_ = 22.02; *p* < 0.001; *η^2^p* = 0.43; see Fig. 3B). There were no differences between groups (rTMS vs. sham-rTMS; PRE: *p* = 0.16; POST: *p* = 0.56).

**Figure 3.**
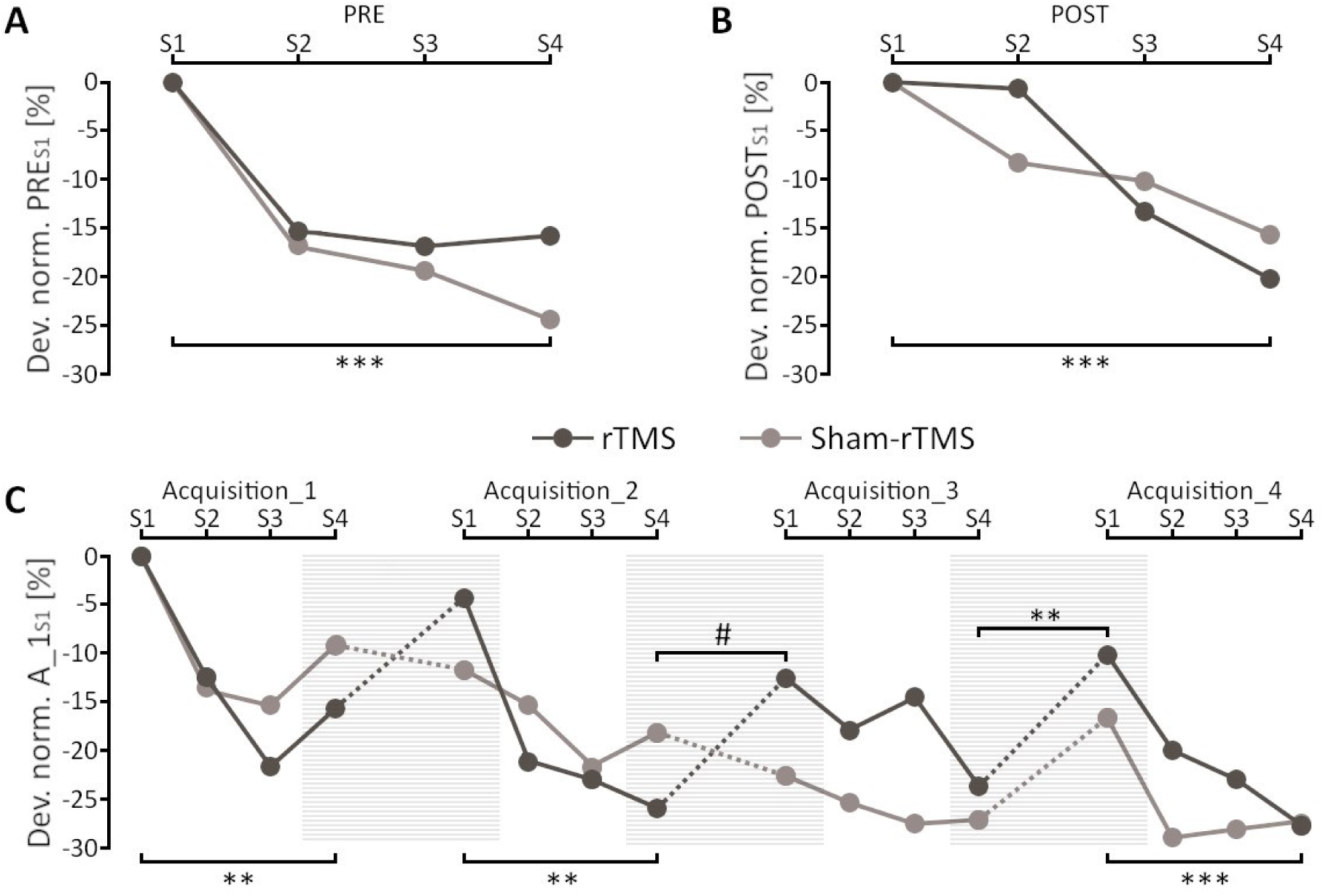
Behavioral data of the balance rocker board task. (**A**) PRE short-term balance acquisition normalized to the first PRE series (PRE_S1_). (**B**) POST short-term balance acquisition normalized to the first POST series (POST_S1_). (**C**) Long-term balance acquisition (solid lines) and consolidation (dotted lines) normalized to the first series of the first acquisition session (A_1_S1_). Shaded areas indicate consolidation after rTMS or sham-rTMS. Dev. norm. = Normalized mean deviation values of the rocker board task. rTMS = repetitive transcranial magnetic stimulation. A_1_S1_ = Acquisition_1, Series 1. *\*\*\** = significant main effect of TIME (*p* < 0.001). ** = significant main effect of TIME (*p* < 0.01). # = significant TIME*GROUP interaction (*p* < 0.01).

Long-term acquisition: Balance acquisition over 4 training sessions (A_1_S1_ to A_4_S4_: *F*_3.73,104.56_ = 8.09; *p* < 0.001; *η^2^p* = 0.22; see Fig. 3C) was highly significant independent of group assignment (rTMS vs. sham-rTMS; A_1_S1_ to A_4_S4_: *p* = 0.64). Thus, both groups had similar performance improvements from A_1_S1_ to A_4_S4_ and there was consequently no significant TIME*GROUP interaction (*p* = 0.30). The analysis of the four individual training-sessions revealed similar performance improvements for both groups, as no significant TIME*GROUP interactions (A_1: *p* = 0.44; A_2: *p* = 0.07; A_3: *p* = 0.44; A_4: *p* = 0.26) or main effects of GROUP occurred (A_1: *p* = 0.44; A_2: *p* = 0.99; A_3: *p* = 0.49; A_4: *p* = 0.60). Significant main effects of TIME were found in three (A_1: *F*_1,29_ = 8.97; *p* = 0.006; *η^2^p* = 0.24, A_2: *F*_1,29_ = 12.80; *p* = 0.001; *η^2^p* = 0.31, A_4: *F*_1,29_ = 22.43; *p* < 0.001; *η^2^p* = 0.45) out of four acquisition sessions and in one session (A_3: *p* = 0.072) the main effect of TIME remained just above the significance level.

### Balance consolidation

The unpaired *t*-test across all consolidation phases together showed a significant difference between the two groups (*t*_29_ = 3.18; *p* = 0.003; *d =* 1.14). The rTMS group displayed a significant performance loss (19 %) compared to the sham-rTMS group, which showed performance stabilization (see fig. 4). More specifically, although the ANOVA from timepoint A_1_S4_ to A_2_S1_ revealed no significant main effects (TIME: *p* = 0.40; GROUP: *p* = 0.96) or TIME*GROUP interaction (*p* = 0.18), the other time points showed significant changes. From timepoint A_2_S4_ to A_3_S1_ the ANOVA revealed a significant interaction of TIME*GROUP (*F*_1,29_ = 11.83; *p* = 0.002; *η^2^p* = 0.29), which indicates differences in consolidation between the rTMS and sham-rTMS group. Tuckey corrected post-hoc tests showed a significant decrease in performance for the rTMS group (*t*_1,29_ = −3.61; *p* = 0.006) and a non-significant increase in performance for the sham-rTMS group (*t*_1,29_ = 1.21; *p* = 0.63). The main effects of GROUP (*p* = 0.90) and TIME (*p* = 0.10) were not significant. The third independent ANOVA for the timepoints A_3_S4_ to A_4_S1_ showed a significant main effect of TIME (*F_1,29_* = 7.84; *p* = 0.009; *η^2^p* = 0.21), no significant effect of GROUP (*p* = 0.64) or TIME*GROUP interaction (*p* = 0.73). However, post-hoc tests revealed significant changes only for the rTMS group (p = 0.036).

**Figure 4.**
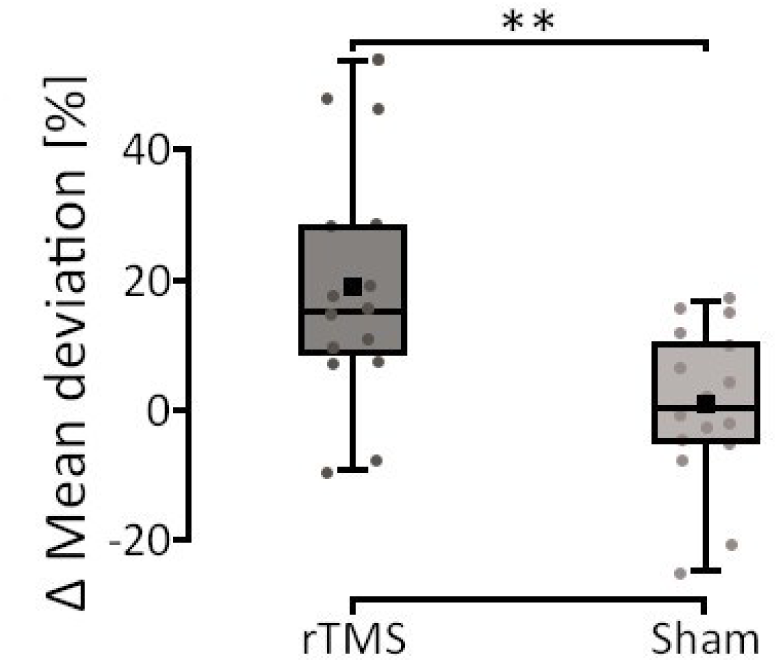
Overview of all three consolidation phases combined, depicting average individual losses or offline gains of each participant, divided by group.

### Short- and long-term effects of rTMS on SICI

Changes in SICI measured during the balancing task on the rocker board were analyzed relative to the first measurement (PRE_1). For all these changes over time, there was a significant main effect of GROUP (*F*_1,29_ = 7.82; *p* = 0.009; *η^2^p* = 0.21; see Fig. 5) indicating that adaptations in SICI differed between the rTMS and the sham-rTMS group. Independent *t*-tests revealed less inhibition for the rTMS compared to the sham-rTMS group for all time comparisons (Δ PRE_1 to PRE_2: *U*_29_ = 58; *p* = 0.014; Δ PRE_1 to POST_1: *t*_29_ = −2.18; *p* = 0.038; Δ PRE_1 to POST_2: *t_29_* = −2.61; *p* = 0.014). The differences between the rTMS and the sham-rTMS group were similar in the short-term (PRE_2) and the long-term (POST_1) as well as after reactivation of the learned task (POST_2).

**Figure 5.**
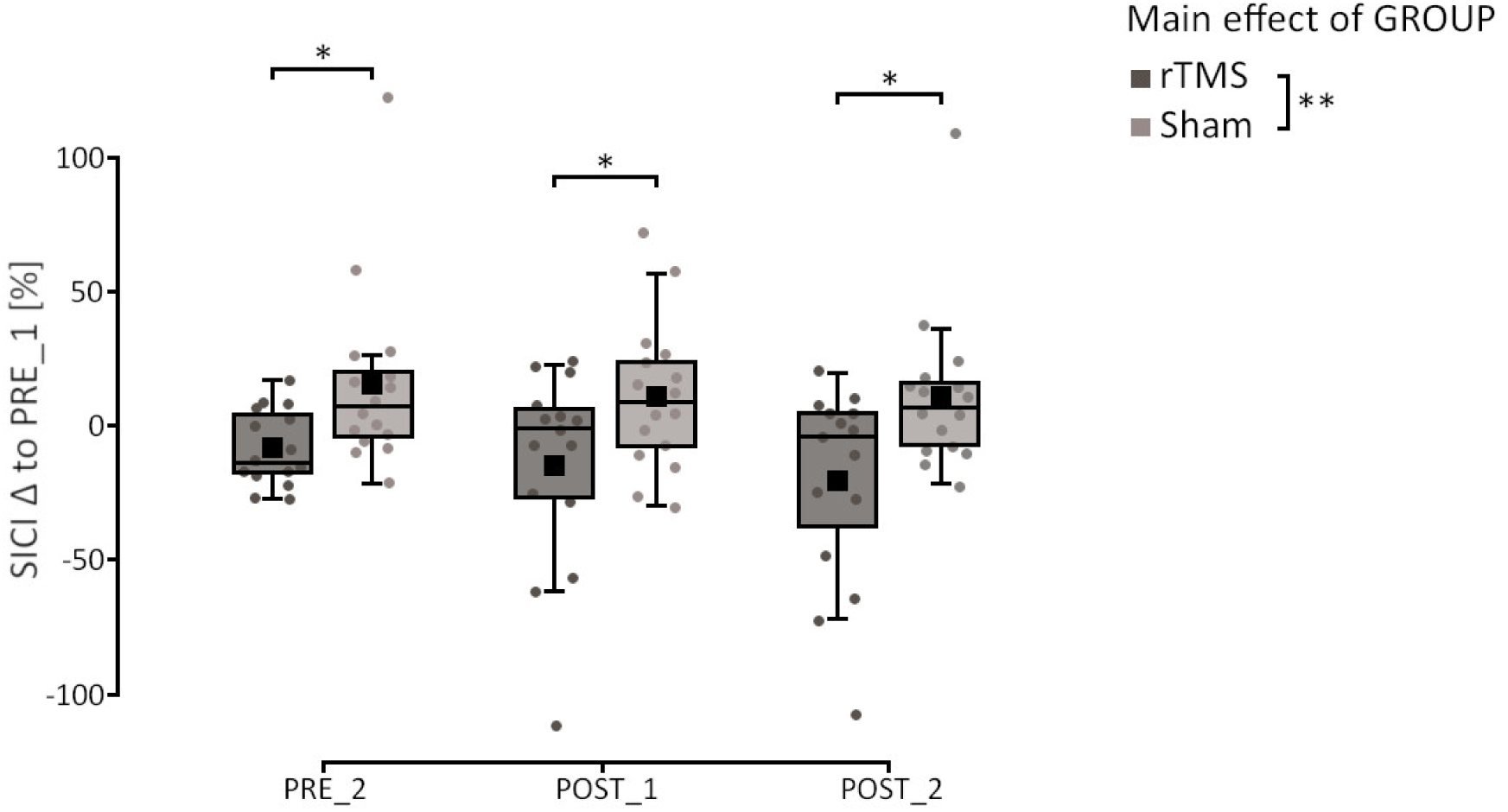
Changes in short-interval intracortical inhibition (SICI) during the balancing condition compared to PRE_1. SICI was up-regulated for the sham-rTMS group in the short-term (PRE_2) and the long-term (POST_1) as well as after reactivation of the learned task (POST_2), whereas SICI in the rTMS group did not increase and was always significantly lower than in the sham-rTMS group at all time points. rTMS = repetitive transcranial magnetic stimulation. Δ = Difference. ** = *p* < 0.01. * = *p* < 0.05.

In contrast, the ANOVA for SICI measurements during the control task (i.e., quiet upright stance) revealed neither significant main effects for TIME (*p* = 0.23) and GROUP (*p* = 0.11) nor any interaction effect (TIME*GROUP: *p* = 0.75) indicating that the adaptations in SICI were specific for the learned task.

### Short- and long-term effects of rTMS on CSE

For the CSE measurements during the balancing task condition, no significant main effects (TIME: *p* = 0.55; GROUP: *p* = 0.63) or interaction (TIME*GROUP: *p* = 0.18) were observed. The same applies to the control condition (quiet stance) with neither significant main effects (TIME: *p* = 0.38; GROUP: *p* = 0.95) nor a significant interaction (TIME*GROUP: *p* = 0.68).

## Discussion

The aim of the present study was to investigate the impact of rTMS on GABA-mediated short-interval intracortical inhibition (SICI) and consolidation of balance performance in the longer-term (i.e., over several acquisition sessions). As hypothesized in the introduction and in line with existing literature (Mouthon & Taube, 2019; Taube et al., 2020), the sham-rTMS group up-regulated SICI after long-term balance acquisition. Moreover, this up-regulation was detectable already in the short-term, i.e., after one balance acquisition session. Conversely, the rTMS group did not display such up-regulation, suggesting that rTMS prevented the modulation of intracortical inhibition. At the same time, rTMS impaired motor memory consolidation of the balance task which was in strong contrast to the sham-rTMS group that tended to have offline gains.

### rTMS-induced interference

For the balance task used in this study (i.e., balancing on the rocker board), we have previously shown short-term interference effects: In our first study, the results indicated that the rocker board task is susceptible to interference when a second balance task is learned shortly afterwards. Based on control experiments, it may even be more specifically said that the two balance tasks should have the same characteristics, such as being controlled by the same muscles, requiring similar control of the center of gravity, and being performed consecutively in order to elicit interference (Egger et al., 2021). In our second short-term study, we used a study design very similar to the current study. More specifically, two groups were asked to train the rocker board task in one single session, after which one group received rTMS and the other sham-rTMS over M1. In the retention test 24 hours later, the rTMS group showed performance losses while the sham-rTMS group showed off-line gains. This indicated that the balance task is susceptible to focally induced interference in M1 in the short-term (Egger et al., 2023). The current results suggest that interference after rTMS over M1 is not only detectable in the short-term but is also present after several learning sessions. Interestingly, the rTMS group showed performance losses across all three consolidation phases, indicating that rTMS had decremental effects even after several learning sessions.

For consolidation processes of complex tasks, it is assumed that other cortical and subcortical motor areas beyond M1 are implicated (Doyon et al., 2009; Doyon & Benali, 2005; Seidler, 2010). As training progresses and motor skills become more automatized, subcortical areas are presumed to play an increasingly pivotal role in motor memory consolidation (Puttemans et al., 2005; Taube et al., 2008). However, based on our results we propose that within the span of the four acquisition sessions, the transition of movement control from more cortical to more subcortical areas has not occurred, yet. This supposition is supported by the consistent performance losses observed in the rTMS group from the end of one acquisition phase to the start of the subsequent phase. This was in strong contrast to the sham-rTMS group, which stabilized performance. Only after the third acquisition phase, a decline in performance was apparent in the sham-rTMS group, too. This decline may be attributed to the possibility that the task difficulty did not reach a sufficient level to elicit further offline gains, potentially indicative of a ceiling effect (Luft & Buitrago, 2005). Consequently, task difficulty emerges as a critical factor in the process of consolidation, with implications for performance outcomes (Akizuki & Ohashi, 2015). Alternatively, this performance decrease may be considered as an outlier, as the balance performance surpassed that of the entire A_3 block.

Notably, both the rTMS and sham-rTMS groups demonstrated comparable performance progression during the acquisition phases, aligning with previous findings indicating that rTMS does not impair acquisition of motor tasks (Muellbacher et al., 2002). This finding is in line with the proposition that rTMS mainly interacts with motor memory consolidation, thereby modulating synaptic plasticity (Hoogendam et al., 2010). The potential for the reversal of synaptic plasticity in the motor cortex, specifically de-potentiation, has been previously documented in humans (Huang et al., 2010). In summary, our results therefore support the assumption that the effects of rTMS have no effect on motor acquisition but interfere with motor consolidation (Censor & Cohen, 2011; Hoogendam et al., 2010), even over multiple acquisition sessions.

### Short- and long-term effects of rTMS on SICI

Previously, a systematic review with meta-analysis about age-related changes in intracortical inhibition after upper extremity learning concluded that there are subtle changes in the brain’s ability to adapt through practice with age and that the age-related decline in SICI does not correlate with the acquisition of motor skills in both healthy young and older adults (Berghuis et al., 2017). However, caution is warranted in interpreting these findings due to certain methodological considerations. Primarily, most studies included in this meta-analysis focused on the effects of short-term learning interventions. This may introduce a notable bias, as it is acknowledged that during the initial learning phase, inhibitory processes are down-regulated to facilitate neural plasticity such as long-term potentiation (LTP). Animal models have demonstrated that transient suppression of local gamma-aminobutyric acid type A (GABA-A) inhibition plays a pivotal role in inducing LTP-like processes, thereby suggesting that reduced GABA release facilitates alpha-amino-3-hydroxy-5-methyl-4- isoxazolepropionic acid receptor-mediated (AMPA) plasticity, crucial for early motor learning stages (Kida & Mitsushima, 2018). Moreover, during early plasticity stages within M1, decreased GABAergic inhibition may facilitate the “unmasking” of pre-existing horizontal connections, thus reinforcing swift remodeling of motor representations (Huntley, 1997; Sanes & Donoghue, 2000). Consequently, the observed general decrease in SICI during short-term studies exploring intracortical inhibitory control in the context of motor learning is not surprising. Secondly, the majority of studies synthesized in Berghuis, and colleagues’ meta-analysis (2017) assessed intracortical inhibition at rest, despite evidence indicating very task-specific modulation of SICI. Recent research has underscored the necessity of evaluating SICI during motor activity, as its modulation varies based on task demands (Opie & Semmler, 2016; Soto et al., 2006). Consistent with this, our recent findings revealed significant up-regulation of SICI during coordinative balance tasks but not at rest or when assessed during ballistic contractions within the same muscle group (Taube et al., 2020). Hence, incorporating motor activity-based assessment of SICI is imperative for a comprehensive understanding of intracortical inhibition dynamics in motor learning paradigms. Therefore, it is more reasonable to assume that SICI is specifically adapted according to the duration of the learning and the motor task itself. For instance, it was consistently shown that SICI will be reduced after strength training (Kidgell et al., 2017; Siddique et al., 2020) or when ballistic contractions are required (to ensure maximal corticospinal output), whereas smooth and coordinative movements such as balancing were accompanied by higher inhibition; probably to allow fine-tuned coordination between different muscles (Mouthon & Taube, 2019; Taube et al., 2020). Interestingly, studies measuring adaptations in SICI during the learned task and in control conditions (at rest or during different tasks) indicated very task-specific adaptations of SICI (Mouthon & Taube, 2019; Taube et al., 2020). In conclusion, cortical inhibition in M1 is likely decreased during acquisition (i.e., initial learning) to promote cortical reorganization (Cirillo et al., 2020; Sanes & Donoghue, 2000) but at later stages can be up- or down-regulated depending on the task (Taube et al., 2020). Furthermore, increases of intracortical inhibition are assumed to promote motor memory consolidation (Cirillo et al., 2020). The results of the current study in the sham-rTMS group are in line with previous observations (Mouthon & Taube, 2019; Taube et al., 2020), indicating increases in SICI after several sessions of balance learning (see fig. 5, POST_1). However, it has to be noted that this is the first study measuring SICI also directly after one single session of balance acquisition. Interestingly and not expected, the amount of SICI increased already in this early phase of learning (see fig. 5; PRE_2). We were surprised to see increases in SICI in the short-term based on the previously mentioned studies in animals, showing transient suppression of GABA for inducing early plasticity such as LTP-like processes and unmasking of horizontal connections (Bachtiar & Stagg, 2014; Huntley, 1997; Sanes & Donoghue, 2000). However, other research has indicated that inhibitory mechanisms can be modulated within minutes (Eisenstein et al., 2023). In particular, this study has shown that a strong inhibitory response immediately after performing the reconsolidation of the task is associated with behavioral changes but that this inhibitory relationship weakens again after some minutes. This suggests that inhibitory mechanisms can be quickly modulated over time according to the requirements for consolidation. As can be clearly seen in fig. 5, rTMS prevented up-regulation of SICI in the short- (PRE_2) and long-term (POST_1) as well as after re-activating the balance task (POST_2). As motor memory consolidation was impaired in the rTMS-group, indicated by high off-line losses between acquisition phases (see fig. 3C, black dots), we assume that the effects of rTMS onto the GABAergic system are an important factor to explain impaired motor memory consolidation after rTMS. This would be in line with studies experimentally manipulating the GABAergic system with drugs such as benzodiazepines (BZD) prior to learning (Bütefisch et al., 2000; Ziemann et al., 2001). BZD serves as positive modulator at the GABA_A_ receptor, increasing the influence of GABA at the receptor. Consequently, these medications are known to inhibit the initiation of LTP, thereby hindering the formation of new memories. Another intriguing effect of BZD emerges when they are administered post-practice, leading to performance enhancements indicating improved motor memory consolidation (Wixted, 2004). It was speculated that recently acquired memories consolidate better after BDZ intake because the formation of new memories is inhibited. If rTMS prevents up-regulation of intracortical inhibition, it can be assumed that the protective effect for the previously acquired task no longer applies. Therefore, our study proposes that rTMS has a down-regulating influence on the GABAergic system (i.e. SICI) which negatively influences motor memory consolidation (Fig. 6).

**Figure 6.**
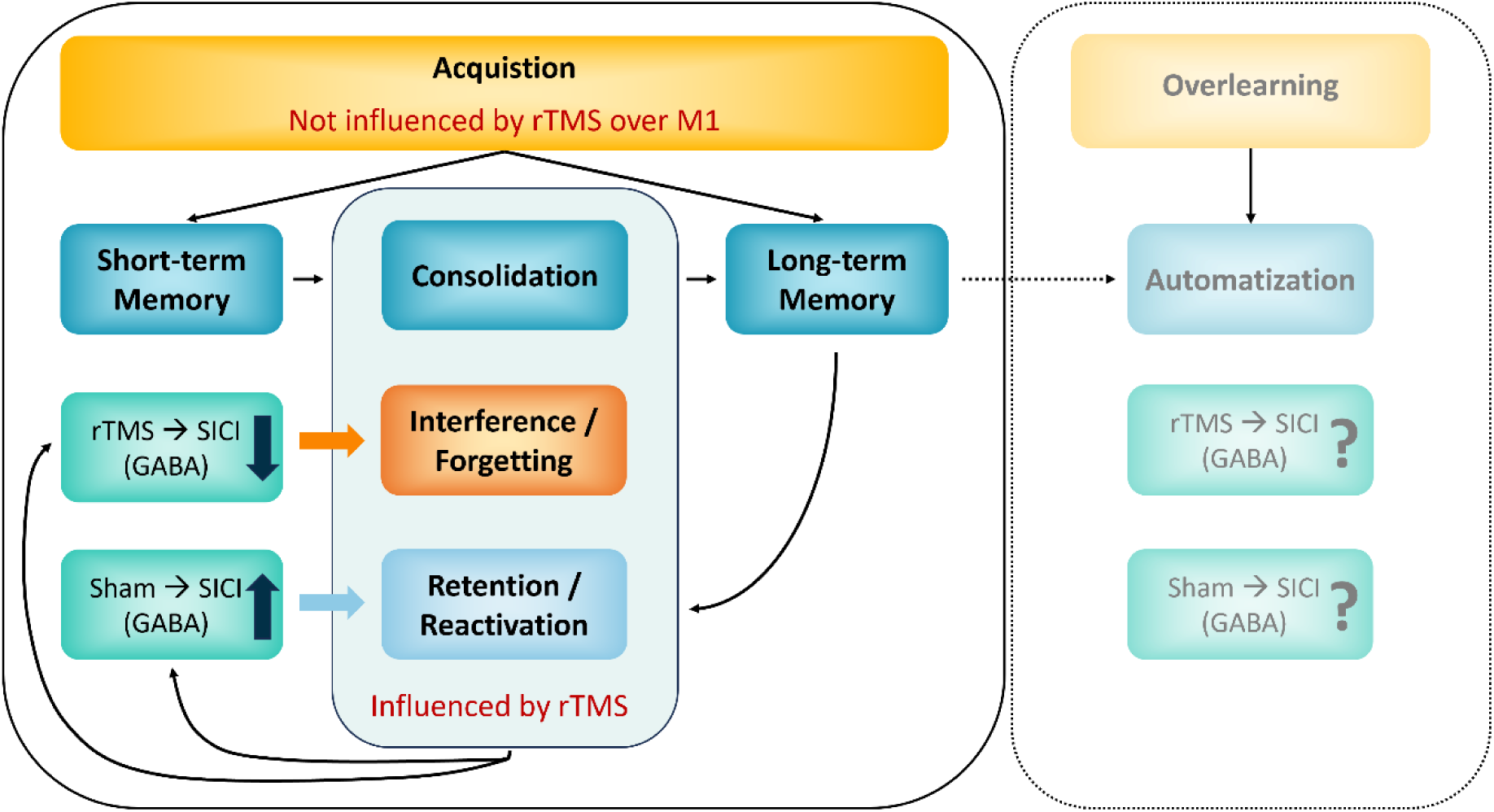
Influence of rTMS on the consolidation process of motor memory. Acquiring a motor task leads to a fragile short-term memory that stabilizes over time, enabling long-term storage and retention of the motor memory. However, applying rTMS during this consolidation phase can interfere with long-term memory, leading to forgetting. These interfering effects might be explained by the suppression of SICI up-regulation by rTMS over M1. The present study did not investigate the long(er)-term effects of balance learning (right, lightly coloured box). However, we would expect that with increasing task automatization (i.e. overlearning), the influence of rTMS on motor memory consolidation and on modulation of SICI are reduced as we would assume a shift in movement control from more cortical to more subcortical areas. SICI = short-interval intracortical inhibition. rTMS = repetitive transcranial magnetic stimulation. M1 = primary motor cortex.

## Conclusion

In summary, although it was known that rTMS impairs the consolidation of motor tasks, the specific mechanisms underlying this interference were not well understood. The present study suggests that rTMS over M1 interferes not only with the consolidation of balance skills but also with up-regulation of SICI (Fig. 6). Therefore, it is proposed to consider modulation of GABAergic mechanisms (i.e., intracortical inhibition) as an important factor for the protection and consolidation of newly acquired motor memories.

## Author contributions

All authors reviewed the manuscript and approved the final version of the manuscript. S.E., M.W. and W.T. concepted the study. S.E., S.M. and M.W. collected the data. S.E. analysed and illustrated the data. S.E. wrote the draft of the manuscript. M.W. and W.T. revised the work.

## Data availability statement

The data that support the findings of this study are available from the corresponding author upon reasonable request.

